# Behavioral tinnitus in gerbils is associated with reduced spontaneous rates in single auditory-nerve fibers

**DOI:** 10.64898/2025.12.01.691754

**Authors:** Amarins N. Heeringa

## Abstract

Tinnitus is often initiated by damage to the peripheral auditory system, for example by overexposure to loud sounds. Animal studies have shown that such noise-induced tinnitus is related to increased spontaneous activity in the brainstem dorsal cochlear nucleus as well as further along the central auditory pathway. However, the role of spontaneous activity of the auditory nerve, connecting the peripheral and central auditory systems, in tinnitus and brainstem hyperactivity remains unknown. In the current study, tinnitus was induced by exposing anesthetized Mongolian gerbils to a 115-dB SPL narrowband noise. After one day of recovery, animals were behaviorally tested for gap detection deficits using a gap-prepulse inhibition of the acoustic startle reflex (GPIAS) paradigm, indicative of tinnitus. Ipsilateral noise-induced threshold shifts did not differ between animals with and without signs of tinnitus. Interestingly, single auditory nerve fibers that were recorded from animals with signs of tinnitus had significantly reduced spontaneous rates compared to both noise-exposed animals without signs of tinnitus and sham-exposed animals. Furthermore, spontaneous rate reduction was specific to fibers with a best frequency within the frequency bands that showed gap detection deficits. On the other hand, inter-spike interval variability and bursting behavior increased in fibers from noise-exposed compared to sham-exposed animals but did not differ with gap detection deficits. These findings suggest that tinnitus-related central hyperactivity may be provoked by reduced spontaneous rates of its innervating auditory nerve fibers. This is consistent with current models explaining the central manifestation of tinnitus, such as the stochastic resonance and central gain models, and allows for a more detailed definition of specifically tinnitus-related deafferentation.

## Introduction

Tinnitus, also known as ‘ringing in the ears’, is defined by the perception of a tonal or noisy sound for which there is no identifiable corresponding external sound source. Prevalence of tinnitus is estimated at 14.4% of adults world-wide who have experienced any tinnitus [1]. In some cases, tinnitus can become extremely bothersome, decreasing quality of life through its impact on sleep, concentration, and mood [2]. Noise exposure is the main risk factor for tinnitus development [3, 4]. This is further illustrated by the finding that 89.5% of young adults report experiencing transient tinnitus after being exposed to loud music [5]. Furthermore, the large majority of tinnitus patients has an abnormal audiogram [6] and those with clinically normal hearing show significantly elevated high-frequency (≥ 10 kHz) hearing thresholds [7, 8]. However, while it is clear that noise-induced cochlear damage is a major risk factor for tinnitus development, it is unclear why some people do, and some do not develop tinnitus after noise-induced hearing loss. Animal studies, where neurophysiological and molecular mechanisms can be assessed in detail, can help solve this important question.

Studies in noise-exposed animals with signs of tinnitus – as determined by established behavioral paradigms – have shown that tinnitus is associated with increased spontaneous firing rates at various stages along the central auditory pathway [9, 10], with the most peripheral stage being the dorsal cochlear nucleus (DCN) [11–13]. Such a tinnitus-related increase in DCN spontaneous rate has been correlated to changes in potassium channel activity that regulate the neuron’s excitability [11, 14] and to altered auditory-somatosensory integration in the DCN principal cells [15–17]. There are several theories available to explain tinnitus-related changes in channel expression, connectivity, and spontaneous activity in the DCN. First, a loss of input, either by degeneration of the auditory nerve (AN) or by hearing loss, can result in a maladaptive increased central gain that amplifies both the sound-evoked and spontaneous activity [18, 19]. Specifically, the concept of stochastic resonance, an adaptive mechanism that reduces the system’s threshold by adding random noise upon a loss of input, has been used to explain the increased spontaneous rates in the DCN [20]. Second, a failure to compensate for a specific loss of input can also explain DCN hyperactivity. This theory poses that DCN hyperactivity is due to the loss of a specific set of neurons with high spontaneous rates – the fast auditory fibers – which in turn reduces the drive for tonic inhibition in the central auditory circuits [21, 22].

It remains under debate whether tinnitus develops due to a unique pathology in the principal cells of the DCN or if tinnitus-specific changes are already apparent at the level of the peripheral auditory system and the AN which in turn trigger DCN pathology. Therefore, the current study aimed at recording from single AN fibers in noise-exposed animals that showed behavioral signs of tinnitus and compared these to AN recordings from noise-exposed animals without behavioral signs of tinnitus. Furthermore, a sham-exposed group allowed to determine the changes that were specific to the noise exposure. As tinnitus and DCN hyperactivity is apparent in the absence of any acoustic stimulation, this study focused specifically on the spontaneous activity of AN fibers. These recordings revealed that noise-exposed animals with behavioral signs of tinnitus had significantly lower spontaneous rates, especially in the fibers that encoded frequencies close the putative tinnitus frequencies, indicated by frequencies that elicited behavioral gap detection deficits. As the recordings were collected shortly after noise exposure, these results suggests that DCN hyperactivity and tinnitus emergence is triggered by a lower spontaneous activity in the auditory nerve. This study can help further explain why, after noise-induced cochlear damage, tinnitus emerges in some subjects but not in others.

## Results

### The gerbil model of tinnitus allows for behavioral separation of noise-exposed gerbils with and without behavioral signs of tinnitus

In the current study, anesthetized Mongolian gerbils were exposed to a 4-kHz centered one-octave narrow-band noise at 115 dB SPL for 75 minutes. The noise exposure was directed unilaterally, to ensure sufficient residual hearing for running the behavioral experiment shortly after. The noise exposure resulted in a significant increase in ipsilateral hearing thresholds, as measured by the auditory brainstem response (ABR), across all tested frequencies immediately after the exposure, which differed per frequency as shown by a significant interaction between time point (pre exposure vs immediately post) and frequency (two-way ANOVA, effect of time point, F(1) = 579.72, p = 2.29*10^-49^, effect of frequency, F(3) = 16.44, p = 4.25*10^-9^, time point x frequency, F(3) = 28.27, p = 4.30*10^-14^; Fig. 1A). The largest threshold elevation immediately after noise exposure was observed at 8 kHz (42.6 dB average threshold elevation), one octave above the center frequency of the narrow-band noise. Three days following noise exposure, thresholds recovered to a large extent but were still significantly elevated compared to baseline values (effect of time point [pre exposure vs 3 days post], F(1) = 131.29, p = 8.61*10^-21^, effect of frequency, F(3) = 14.14, p = 6.49*10^-8^, time point x frequency, F(3) = 10.63, p = 3.15*10^-6^; Fig. 1A). Sham exposure, where a different set of animals were submitted to the same paradigm except with the speaker disconnected, did not result in significant threshold changes, neither immediately (effect of time point, F(1) = 0.50, p = 0.49) nor 3 days after the sham exposure (F(1) = 0.12, p = 0.74; Fig. 1B).

**Figure 1.**
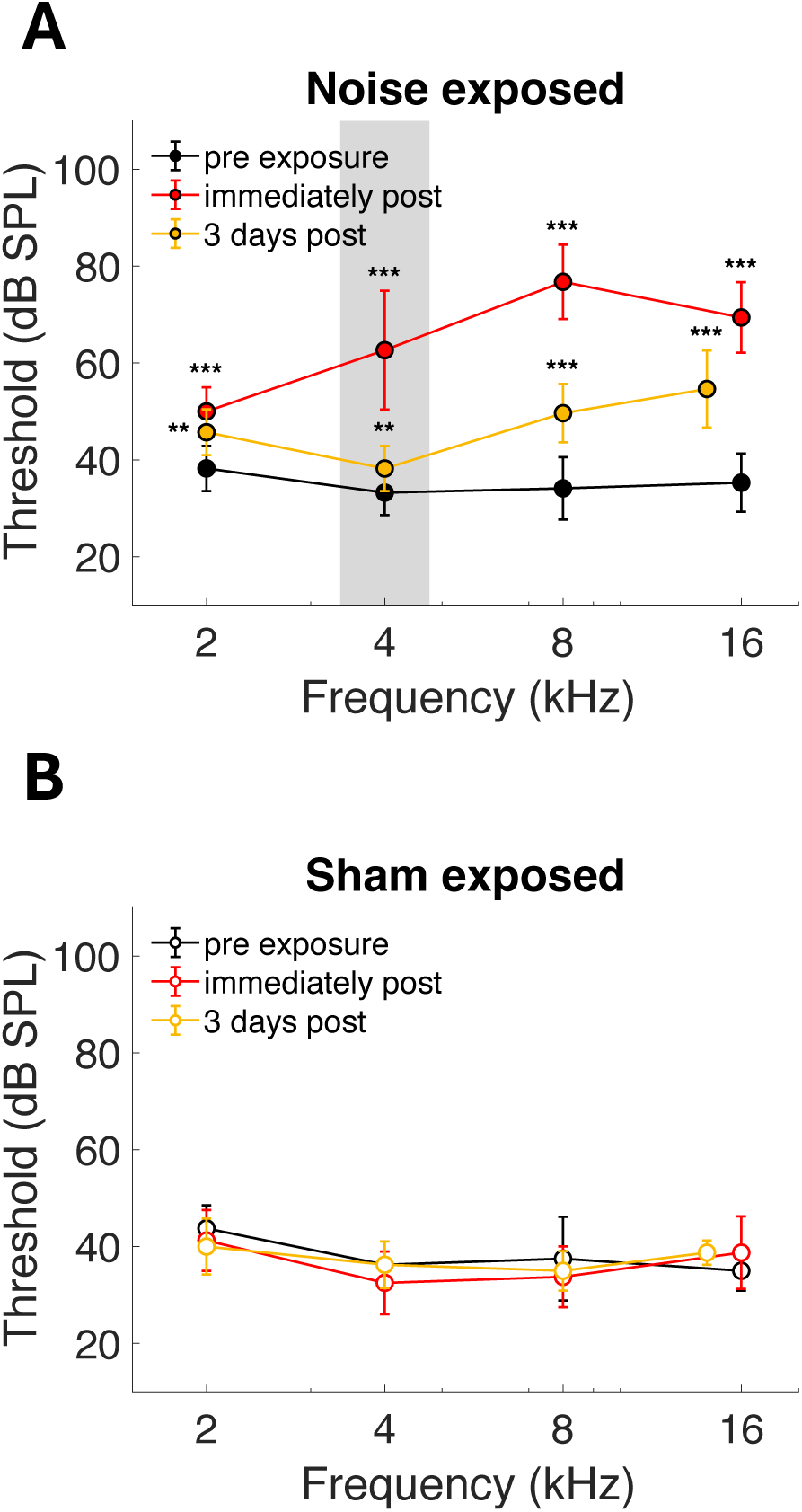
The effect of noise- and sham exposure on ABR hearing thresholds. A) ABR thresholds of noise-exposed gerbils (n = 15) before the noise exposure (black markers), immediately after the noise exposure (red markers), and 3 days after the noise exposure on the day of recording from single-unit auditory nerve fibers (yellow markers). B) The same plot as in panel A, but for the gerbils that were sham exposed (n = 4). Data are shown as mean ± STD. ** indicates p < 0.01 and *** indicates p < 0.001 following post-hoc tests. P-values were corrected for multiple comparisons. The grey area indicates the frequency region of the noise exposure.

To test whether a noise-exposed animal showed signs of tinnitus, the Open-(G)PIAS software by Gerum, Rahlfs (23) was implemented in a set up that measures the gerbil’s acoustic startle reflex. Typically, the amplitude of startle reflexes were larger in trials without a gap compared to trials with a gap preceding the startle stimulus (see Fig. 2A for an example). Gap-PPI is calculated as 1 minus the ratio of the startle amplitude in gap trials over the startle amplitude in no-gap trials and is expressed in percentages, with lower values indicating less inhibition of the acoustic startle response by the presence of the gap, and thus presumably worse gap detection. When gap-PPI was significantly reduced post exposure at any frequency for a given animal, the animal was classified with behavioral signs of tinnitus (TI animal, see Fig. 2B for an example). GPIAS was assessed two days after the noise exposure, to allow the animal to recover from the exposure and the anesthesia. Out of the 15 noise-exposed animals, 5 were classified with signs of tinnitus. The frequency band with significant gap detection deficits was not specific to the center frequency of the noise exposure and differed per individual, consistent with previous reports showing that gap detection deficits become more specific to the exposure frequency over longer periods of time [24, 25]. Animals with no signs of tinnitus (NT animals) have no significant decreases in gap-PPI by definition (see Fig. 2C for an example) and sham-exposed animals also did not show any gap detection deficits (see Fig. 2D for an example).

**Figure 2.**
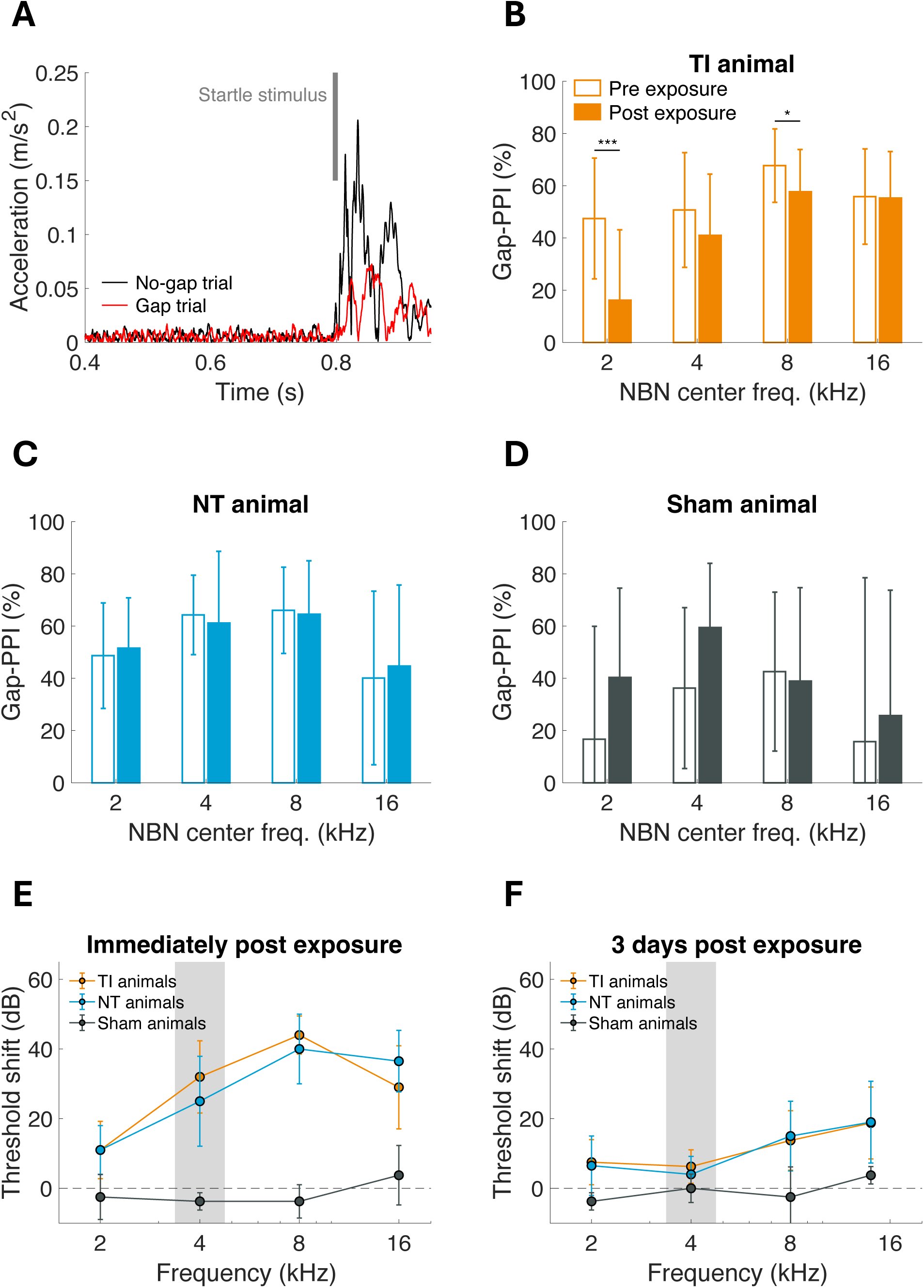
GPIAS allows for behavioral separation of animals with and without signs of tinnitus. A) An example trace of a startle response in a no-gap trial (black trace) and a gap trial (red trace). The time of the startle stimulus, which is at 0.8 s after the trigger that starts collecting the acceleration data, is indicated in grey. Startle amplitude is determined by the maximum acceleration following the startle stimulus. Gap-prepulse inhibition (gap-PPI), the relative difference between the two startle amplitudes, is 64% for this combination of trials, i.e. there is a 64% reduction in startle amplitude due to the presence of the gap. Data derived from a baseline session of a random animal at a 4-kHz center frequency narrowband noise (NBN). B) Gap-PPI (in %) across four different center frequencies of the background NBN from an example animal with significant gap detection deficits after noise exposure (TI animal). Baseline gap-PPI is presented in open bars, gap-PPI after noise exposure is shown in filled bars. * indicates p < 0.05 and *** indicates p < 0.001 following T-tests, corrected for multiple comparisons. C) Gap-PPI of an example animal without post-exposure gap detection deficits (NT animal). Baseline gap-PPI is presented in open bars, gap-PPI after noise exposure is shown in filled bars. D) Gap-PPI of an example sham-exposed animal. Baseline gap-PPI is presented in open bars, gap-PPI after noise exposure is shown in filled bars. E) ABR threshold shifts immediately following the exposure plotted separately for the noise-exposed gerbils with signs of tinnitus (TI, in orange) and those without signs of tinnitus (NT, in blue), and for sham-exposed animals (in black). D) Threshold shifts 3 days after the exposure. Data are shown as mean ± STD. The grey area indicates the frequency region of the noise exposure.

To determine whether behavioral signs of tinnitus were related to the extent of hearing threshold elevations, the ABR data converted to thresholds shifts were plotted separately for the three different experimental groups (Fig. 2E and 2F). No significant differences were observed between threshold shifts of animals with compared to animals without signs of tinnitus at either time point (immediately after exposure: effect of behavioral outcome, F(1) = 0.11, p = 0.74, Fig. 2C; 3 days after exposure: F(1) = 0.03, p = 0.87, Fig. 2D). This suggests that the development of behavioral signs of tinnitus did not depend on the extent of ABR hearing threshold elevations.

### Spontaneous firing rates were reduced in animals with behavioral signs of tinnitus

Single-unit AN fiber recordings were obtained from noise-exposed animals with and without behavioral signs of tinnitus and sham-exposed animals 3 days after the noise exposure. Figure 3A shows the spontaneous rate (SR) of these fibers as a function of their best frequency. Average SR differed significantly among the three experimental groups (Kruskal-Wallis test, χ^2^ = 11.07, p = 0.004; Fig. 3B). Specifically, fibers recorded from animals with behavioral signs of tinnitus (TI) had significantly lower spontaneous rates than those recorded from animals without signs of tinnitus (Wilcoxon Rank Sum test, Z = 3.13, p = 0.0055) and compared to those recorded from sham-exposed animals (Z = 2.87, p = 0.0082). Noise-exposed animals without signs of tinnitus (NT) did not show a significant change in average spontaneous rate when compared to sham-exposed animals (Z = 0.13, p = 0.90).

**Figure 3.**
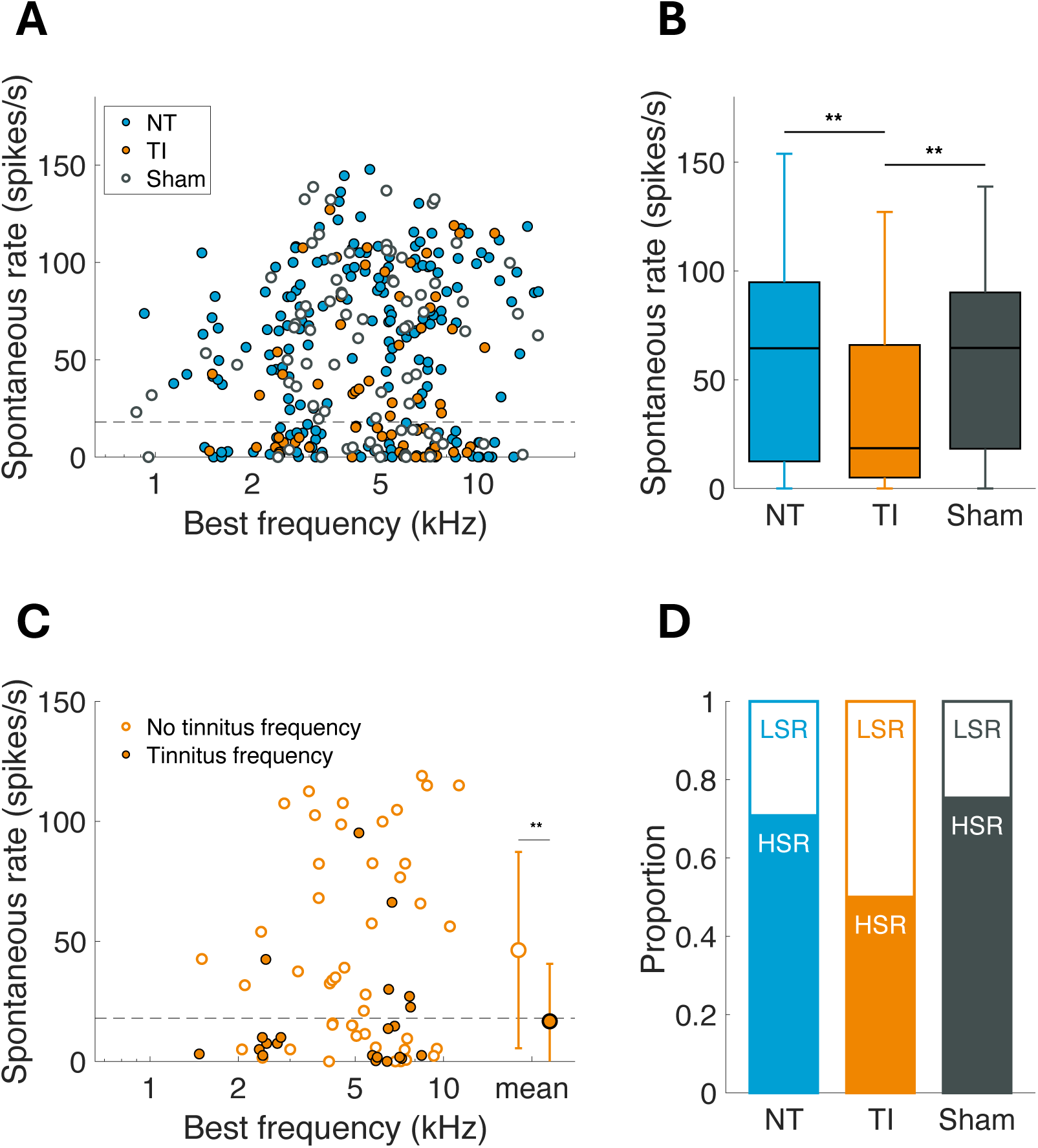
Average spontaneous rate was decreased in auditory nerve fibers recorded from noise-exposed animals with signs of tinnitus. A) Spontaneous rate plotted as a function of best frequency for fibers recorded from sham-exposed animals (open, black markers), noise-exposed animals with no signs of tinnitus (NT; blue markers), and noise-exposed animals with signs of tinnitus (TI; orange markers). The dashed line indicates 18 spikes/s, the threshold for considering a fiber as low- vs high-spontaneous rate [26]. B) Average spontaneous rates for the three experimental groups separately. Boxplots show the median, the 25^th^ and 75^th^ percentile, and the range of data points in error bars. Significant differences are indicated in the plot, ** indicates p < 0.01 following post-hoc tests with p-values corrected for multiple comparisons. C) Data of animals with signs of tinnitus plotted separately for the fibers with a best frequency within the frequency range that showed gap detection deficits (filled markers) and for those outside of this frequency range (open markers). ** indicates p < 0.01. D) Data plotted as proportion of fibers with a low-spontaneous rate (LSR) and with a high-spontaneous rate (HSR) for the three experimental groups.

The GPIAS model also allowed identification of the frequency bands at which the gap detection deficits were significant. Gap detection deficits were observed at narrowband noises centered at 2, 4, and 8 kHz, but differed per animal classified with tinnitus. Fibers recorded from the animals with signs of tinnitus were also grouped according to whether its best frequency fell within the frequency range of a narrowband noise band that yielded gap detection deficits in a given animal. Spontaneous rates of fibers that were within such frequency bands with gap detection deficits were significantly lower compared to those that were not (Z = 3.02, p = 0.0025; Fig. 3C).

AN fibers are typically grouped into those with a low SR and those with a high SR, with a cut-off rate at 18 spikes/s [26], which correlate to both physiological and molecular profiles [27, 28]. The proportion of low-SR fibers in sham-exposed animals was 0.25, which was similar to the proportion of low-SR fibers recorded in naïve young-adult gerbils from previous studies and other labs [29, 30]. In fibers recorded from animals without signs of tinnitus, the distribution of SR classes did not significantly differ from those of sham-exposed animals as determined by a bootstrap analysis (low-SR fibers, 0.29: Z test, p = 0.13; high-SR fibers, 0.71: p = 0.15; Fig. 3D). However, the distribution of SR classes in fibers recorded from animals with signs of tinnitus significantly differed from those recorded in sham-exposed animals, determined by bootstrap analysis (low-SR fibers, 0.50: p = 1.53*10^-6^; high-SR fibers, 0.50: p = 8.78*10^-6^; Fig. 3D), with a proportion of 0.5 low-SR fibers.

Together, these analyses show that fibers recorded from noise-exposed gerbils with signs of tinnitus had a significantly lower spontaneous rate, especially if its BF was close to the frequency that showed gap detection deficits.

### Patterns in spontaneous activity were affected by noise exposure but not by gap detection deficits

Acoustic information is not only transferred to the brain via average rates, but also through the underlying patterns within the spike times. Most of these patterns are determined in relation to the presented sound, such as first-spike latency, jitter, or phase locking. However, there are also some independent measures that can be used to determine the degree of patterning in spontaneous activity. First, the coefficient of variation (CV) was calculated from inter-spike intervals of long spontaneous activity recordings (∼3 min), which reveals the degree of variability in the instantaneous firing rate. For high-SR fibers (> 18 spikes/s), the CV correlates with the SR. An analysis of covariance reveals this strong correlation to SR and a significant difference between the three experimental groups (ANCOVA, SR factor, F(1) = 404.95, p = 4.49*10^-44^, effect of experimental group, F(2) = 5.84, p = 0.0036; Fig. 4A). Post-hoc analyses showed significantly higher CVs (more variability in inter-spike intervals) only for the fibers from animals without signs of tinnitus compared to the sham-exposed group (p = 7.83*10^-4^), when controlling for SR, but not between the other groups. For low-SR fibers, there is more variability in CV between fibers but no statistical difference between the three groups (Kruskal-Wallis test, χ^2^ = 0.95, p = 0.62; Fig. 4A).

**Figure 4.**
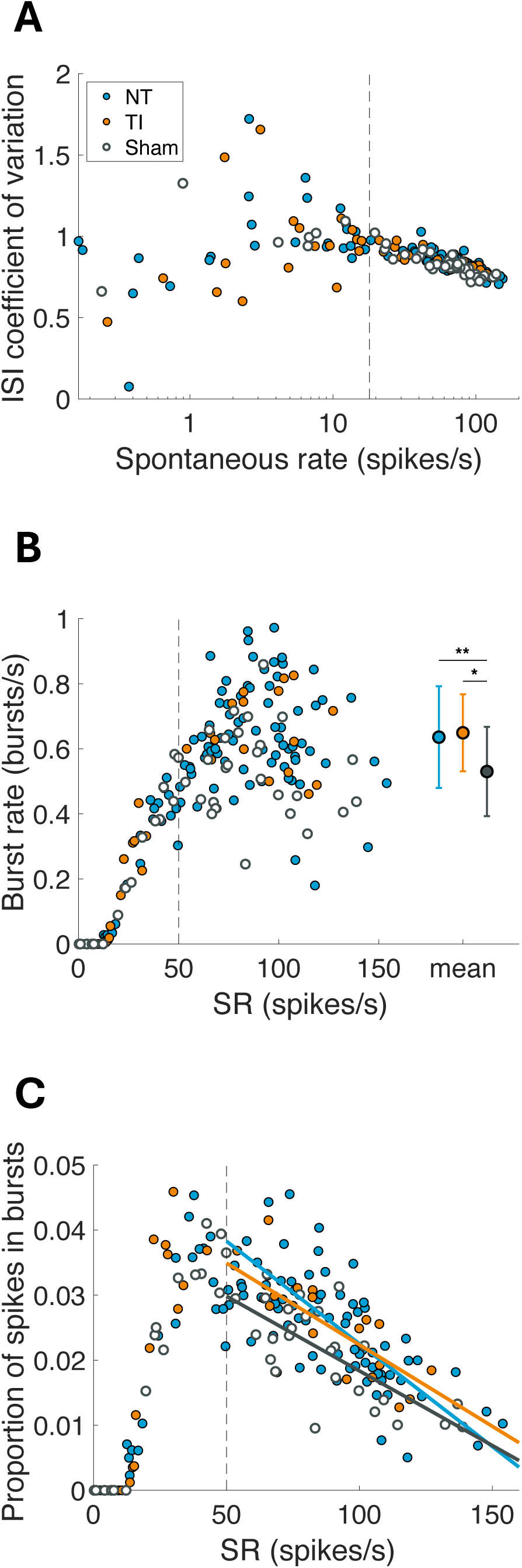
Patterns in spontaneous activity differed between the noise- and sham-exposed groups. A) Coefficient of variation (CV) for the inter-spike intervals (ISI) plotted as a function of spontaneous rate. A high CV indicates more variability between ISIs, in other words higher variability in instantaneous spontaneous rate across a long recording in silence. The dashed line indicates the cut-off for high- vs low-SR fibers, at 18 spikes/s. Noise-exposed animals without gap detection deficits are plotted in blue markers (NT), noise-exposed animals with gap detection deficits are plotted in orange markers (TI), and sham-exposed animals are plotted in open markers (Sham). B) Burst rate, in bursts/s, plotted as a function of spontaneous rate. Note that here SR is plotted on a linear scale to illustrate the differences at high rates. The legend of panel A applies. The dashed line indicates here the cut-off at 50 spikes/s, which indicates stabilization of bursting behavior across SR. The mean ± STD are shown in larger symbols at the right. * indicates p < 0.05, ** indicates p < 0.01 following post-hoc tests, corrected for multiple comparisons. C) Proportion of spikes in bursts plotted as a function of spontaneous rate. The dashed line indicates the cut-off at 50 spikes/s, similar to panel B. The legend of panel A applies. The trend lines show linear regressions, separately for noise-exposed without tinnitus (blue), noise-exposed with tinnitus (orange), and sham-exposed animals (black).

One explanation for increased variability in inter-spike intervals is the enhanced presence of bursting behavior. Bursting was analyzed using the Gaussian Surprise method by Ko, Wilson (31). Bursting behavior, as determined by several burst-related variables, was significantly different between the fibers recorded from both noise-exposed groups (TI and NT animals) compared to those from sham-exposed animals. Figure 4B shows the burst rate, i.e. the average number of spontaneous bursts per second, for the three experimental groups. Only fibers with rates higher than 12 spikes/s showed significant bursting behavior in the spontaneous activity, according to the robust Gaussian surprise method. Burst rate first steeply increased with SR and then stabilized at around 50 spikes/s. Therefore, to statistically test for differences between the groups, only fibers with an SR above 50 spikes/s were included in the comparison, which showed a significant difference between experimental groups (Kruskal-Wallis test, χ^2^ = 10.70, p = 0.0047; Fig. 4B). Post-hoc analyses revealed significantly higher burst rates for the fibers from animals that were noise exposed (NT vs Sham, p = 0.0066; TI vs Sham, p = 0.013), but not between fibers from animals with compared to those without signs of tinnitus (NT vs TI, p = 0.73; Fig. 4B).

Correspondingly, the proportion of spikes that were part of a burst also showed a significant effect of experimental group (ANCOVA, SR factor, F(1) = 129.86, p = 1.52*10^-20^, effect of experimental group, F(2) = 6.45, p = 0.0022; Fig. 4C). Specifically, the fibers from the sham-exposed animals had a lower proportion of spikes in bursts compared to the fibers from both noise-exposed groups when corrected for SR, regardless of the behavioral outcomes of the GPIAS test (NT vs Sham, p = 0.014; TI vs Sham, p = 0.030; NT vs TI, p = 0.85, p-values from ANCOVAs and corrected for multiple comparisons).

Together, these analyses show that fibers recorded from noise-exposed gerbils had higher variability in their instantaneous firing rates and displayed more bursting than fibers from sham-exposed gerbils, while no significant differences were found when comparing between fibers from animals with vs without signs of tinnitus.

## Discussion

The current study examined whether spontaneous activity of single AN fibers was affected by noise-induced gap detection deficits in the GPIAS behavioral model, indicative of signs of tinnitus. Average spontaneous rates were significantly lower in animals with behavioral signs of tinnitus, especially in fibers with a BF within a frequency band of the narrowband noise that showed gap detection deficits. When the inter-spike intervals of the spontaneous activity were studied more closely, significant changes in firing rate variability and in bursting behavior were observed with noise exposure, but not with gap detection deficits. Similarly, auditory thresholds were affected only by noise exposure but did not differ between animals with and without signs of tinnitus. As the recordings were made shortly after the noise exposure (3 days), these results suggest that the emergence of tinnitus-related DCN hyperactivity corresponds to a reduced spontaneous rate of single auditory nerve fibers.

### The relation between reduced AN spontaneous rate and tinnitus

The conclusion that AN spontaneous rate is reduced with tinnitus is based on the important, critical assumption that gap detection deficits of noise-exposed gerbils in the GPIAS behavioral model are an accurate measure for tinnitus in animals. Human studies show conflicting results, with both evidence for gap detection deficits in humans with tinnitus [32, 33], and no difference in gap detection abilities between humans with and without tinnitus [34–36]. Nevertheless, much of what is known about the central pathology of tinnitus derives from studies in animals with gap detection deficits [15, 37, 38]. Even though the results presented here can be well related to these important neurophysiological and histological findings, they should be handled with care when translating to the human condition.

A reduction in AN fiber spontaneous rate following noise exposure is consistent with previous studies. However, both the recovery time and the extent of the threshold shift seem to affect whether and to which extent a reduced spontaneous rate in single AN fibers can be observed. For example, following an intense noise exposure resulting in threshold shifts of at least 20 dB and after recovery times longer than 4 weeks, reduced AN spontaneous rates are evident in both cats and guinea pigs [39, 40]. On the other hand, when the noise-induced threshold shifts are temporary and recovery times are longer than 2 weeks, spontaneous rates remain unchanged in CBA/CaJ mice [41] or show a selective loss of fibers with a low-SR in guinea pigs [42]. When recording from the cochlear round window in silence, which is a proxy for the summed spontaneous activity in the AN, at different time points after noise exposure in gerbils, Jeffers, Bourien (43) found a significant decrease of round window activity 24 h after the exposure, which was recovered 2 weeks after the exposure. These studies together would suggest that SR reductions are mainly correlated to the noise-induced thresholds shifts. However, the current findings showed significant SR reductions in gerbils with signs of tinnitus, but not in those without signs of tinnitus. Importantly, threshold shifts did not differ between the gerbils with and without signs of tinnitus. For both groups, thresholds were improved compared to immediately after the noise exposure at the time of recording from the AN but did fully recover (yet) to baseline thresholds.

There are several possible mechanisms that could cause SR reductions in the AN following noise exposure. From the studies in cats with permanent threshold shifts, it was shown that the SR reductions are associated with a selective loss of the tallest row of stereocilia from the inner-hair cells [39]. Furthermore, noise exposure can also lead to an acute decrease in the endocochlear potential [44], which reduces the AN spontaneous rate as well [45]. A specific loss of high-SR fibers can also theoretically be responsible. However, since low-SR fibers are typically the most vulnerable to any kind of cochlear damage, such as exposure to ototoxic compounds, age-related hearing loss, and noise exposure with a temporary threshold shift [30, 42, 46], this possibility seems improbable.

Tinnitus can also be experimentally induced by administration of sodium salicylate, the active compound of aspirin. Similar to the results presented here, it has been shown in both guinea pigs and Mongolian gerbils that salicylate administration leads to a reduction of spontaneous rate in the auditory nerve [47–49]. While the mechanism causing the spontaneous rate reduction may not be the same between salicylate administration and noise exposure, the central consequences of the phantom perception may be associated to this same phenomenon.

### Implications for tinnitus-related pathology in the dorsal cochlear nucleus

The aim of the current study was to determine whether tinnitus-specific changes are already apparent at the level of the AN which in turn could trigger the DCN pathology. The findings suggest that a reduced auditory nerve fiber spontaneous rate may initiate such changes at the level of the DCN. On the other hand, changes in variability and bursting behavior of the remaining spontaneous activity in the auditory nerve fibers were not associated to behavioral signs of tinnitus, but rather to noise exposure in general. This suggests that the reduction of spontaneous rate, rather than a specific arrangement of the spikes, is associated to central tinnitus pathology. This finding is consistent with the theory that tinnitus is caused by a loss of fibers with high spontaneous rates, the so-called ‘fast auditory fibers’ [21, 22]. By reduced average spontaneous rate and/or a specific loss of high-SR fibers, the drive for tonic inhibition in the central auditory system diminishes and as such may contribute to tinnitus pathology.

The recordings in the current study were collected three days after noise exposure, suggesting a specific role in the early stage of tinnitus pathology. The study by Jeffers, Bourien (43) on round window activity in noise exposed gerbils suggests that spontaneous rates recover again to normal levels two weeks after noise exposure, with a concurrent recovery of auditory thresholds. Whether the reduced spontaneous rate recovers again or remains low in animals with specifically behavioral signs of tinnitus remains to be studied. Furthermore, when recording from single AN fibers, activity can only be measured from those fibers that retained a functional connection to the inner hair cell. Therefore, a specific loss of auditory nerve fibers may have occurred in parallel, putatively also contributing to tinnitus-related central pathology. This question has been studied in both humans and laboratory animals, with variable outcomes. Some studies show that tinnitus is associated with a reduced number of functional synapses between inner hair cells and AN fibers [37, 50–52], whereas others do not find differences between tinnitus and control groups [17, 38, 53].

The finding of a reduction of spontaneous rate in single auditory nerve fibers can help further refine current models of central manifestation of tinnitus, such as the stochastic resonance and central gain models [20, 22]. As such, a more detailed definition of peripheral deafferentation in tinnitus could be offered, that bridges between knowledge on central pathologies and peripheral damage in tinnitus.

## Methods

### Animals

Nineteen Mongolian gerbils (*Meriones unguiculatus*) were used in this study. Animals were of either sex (13 female) and weighted between 49 – 103 g at the final day of the experiment. At the start of the experimental procedures, animals were between 16 – 21 weeks old, i.e. young adult. The full experimental procedure took between 4 and 6 weeks. The first two weeks were for the animals to habituate to the behavioral setup and in the third week, baseline behavioral data was collected. In the following weeks, each animal was noise exposed, behaviorally tested for tinnitus, and subjected to a final experiment where single units were recorded from the auditory nerve. Gerbils were born in the gerbil colony at the animal facility of the University of Oldenburg and were raised and housed in a climate-controlled, quiet environment. Animals were fed *ad libitum*, were provided with cage enrichment, and were housed in a group with three same-sex littermates when possible. Out of the 19 animals, 15 were noise exposed to induce tinnitus, while 4 were sham exposed and served as a control group. The noise exposed animals were further subdivided between those that exhibited behavioral deficits in the GPIAS paradigm (n = 5), suggestive of tinnitus, and those that did not show such deficits (n = 10). Experimental procedures were approved by the ethics authorities of Lower Saxony, Germany (LAVES), under permit number 33.19-42502-04-22-00167.

### Gap prepulse inhibition of the acoustic startle response

#### Experimental setup

To evaluate whether an animal perceived tinnitus, the gap prepulse inhibition of the acoustic startle response (GPIAS) paradigm was employed [54, 55]. In this paradigm, the amplitude of the acoustic startle reflex is evaluated when the startle stimulus is presented in a background noise and when the startle stimulus is preceded by a short gap in the background noise. In normal-hearing animals, the gap typically results in a smaller startle response, termed gap pre-pulse inhibition (gap-PPI) (see Fig. 1A). The paradigm assumes that animals with tinnitus exhibit impaired gap detection, resulting in a smaller degree of gap-PPI.

The experimental setup used in the current study was custom-build based on the setup developed by Gerum, Rahlfs (23). Briefly, the animal was sitting in a rectangular mesh-wired fixation cage (13 x 6 x 6 cm), which allowed for free movement of the animal. The fixation cage was tightly mounted on a light-weight metal plate topped with a 1-cm thick layer of acoustic foam. The metal-plate was standing on four springs (0X-RDF1406, Febrotec), one in each corner, to allow for flexible movement of the fixation cage. An acceleration sensor (ADXL 337 on a GY-61, Conrad) was connected to the bottom of the metal plate to measure startle responses. Three-dimensional acceleration data (x, y, and z dimension) was collected and digitized using a breakout box (BNC-2110, National Instruments; sampling rate 1000 Hz) and stored on a computer using a data acquisition card (PCIe-6320, National Instruments).

Acoustic stimuli were presented using an external audiocard (FireFace UCX, RME; sampling rate 96 kHz), amplified (RMB 1506, Rotel), and delivered through two loudspeakers that were mounted to the ceiling of the acoustic chamber using metal wire. The speakers were positioned 30 cm above the fixation cage that held the experimental animal, to enable a balanced acoustic field irrespective of the positioning of the animal. A Canton speaker (Canton Plus XS.2) was used to deliver the background stimulus and a tweeter (Peerless XT25TG30-04, Quint Audio) was used to deliver the startle stimulus. Speakers were equalized and calibrated using a ½” free-field microphone (type 40AF, GRAS) connected to a pre-amplifier (type 26AK, GRAS) and a power module (type 12AD, GRAS).

Open(G)PIAS, an open-source software program written in Python, controlled the setup by presenting the acoustic stimuli and simultaneously collecting the startle response data [23]. A trigger, send from the soundcard into the breakout box, assured synchronization between acoustic stimulation and data acquisition. The setup was fixed on a low-vibration table (Newport), placed in a sound-proofed chamber (Industrial Acoustic Company, GmbH) that was aligned with two layers of acoustic foam to reduce reverberations (Plano 50 and Pyramid 50; Pinta Acoustics). The animal was monitored during the experiment via a camera.

#### Experimental procedure

To habituate the animals to the setup, first the fixation cage was introduced to the animals by placing it in their home cage and leaving it there for a week. Using sunflower seeds to lead the gerbils into the fixation cage, the gerbils were then placed into the setup for 30 minutes without any acoustic stimulation at two consecutive days. Next, two to three baseline GPIAS sessions were collected from each animal, with at least three days between each baseline session. A GPIAS session consisted of 240 trials in which the startle amplitude was measured in response to a 105 dB SPL white noise burst (20-ms duration). There was a randomized interval between startle stimuli, ranging from 11 to 15 s. During each trial, a bandpass filtered octave-wide narrowband background noise was presented at 60 dB SPL RMS with center frequencies of either 2, 4, 8, or 16 kHz. Each center frequency was presented for 60 trials, in a pseudorandomized order. Furthermore, in half of the trials, a gap (50 ms duration, 2 ms cos^2^ ramps) in the background noise was presented before the startle stimulus with a 100-ms lead time. A 30 – 40-minute break was introduced when half of the trials were presented, for the animal to rest in its home cage. At the start of each session and after the break, five trials without any background noise were presented to allow for adaptation to the startle stimulus. These startle amplitudes were discarded in the analysis. GPIAS was measured again two days after the noise exposure. All GPIAS sessions were performed at similar times during the day for one animal, typically in the afternoon.

#### Data analysis

Raw data traces from the acceleration sensor were low-pass filtered (45 Hz cut-off), multiplied by their calibration factors, and integrated into one acceleration measure by taking the root-mean-square at each time point. The maximum acceleration within a 150-ms time window following startle stimulus onset was considered the startle amplitude. When the acceleration in a 400-ms time window before the startle stimulus exceeded a threshold of 0.1 m/s^2^, the trial was removed from further analysis due to movement artifacts.

Initial visualization of the startle amplitudes (*A*) in gap and no-gap trials for the four different center frequencies was performed using a custom-written Python script (run in Spyder version 5, Python 3.11). Subsequently, gap-PPI was analyzed for baseline and post-noise exposure sessions using custom-written scripts in MATLAB (version R2023b, Mathworks).

*PPI* was calculated for each background frequency in each session according to:

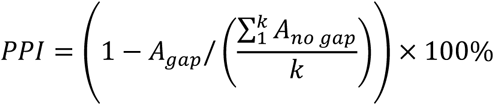

with *A* indicating the startle amplitude and *k* indicating the number of included no-gap trials for one center frequency. *PPI* will thus be calculated for each gap trial in percentage, with lower values indicating poorer gap detection.

### Noise exposure

#### Anesthesia

Animals were anaesthetized with ketamine (135 mg/kg; Ketamin, Serumwerk) and xylazine (6 mg/kg; Xylazin, Serumwerk), which was maintained with one-third of the initial dose hourly or one-sixth of the initial dose when the hindpaw reflex was positive. In addition, all animals received a subcutaneous injection of meloxicam (0.5 mg/kg; Metacam, Boehringer Ingelheim), oxygen flowing onto the snout (1.5 L/min), eye ointment (Bepanthen, Bayer Vital GmbH), and a subcutaneous injection of saline (sterile 0.9% NaCl solution, 4 ml). Six out of nineteen animals also received a prophylactic dose of atropine at the start of the experiment (0.3 mg/kg; Atropinsulfat, B. Braun). Heartbeat and breathing were constantly monitored through an electrocardiogram, which was visualized on an oscilloscope (Voltcraft 630-2) and made audible on a speaker (Go2, JBL). Body temperature was maintained at 38 °C using a homeothermic blanket (Harvard Apparatus).

#### Experimental procedure

Before noise exposure, hearing thresholds were determined using the auditory brainstem response (ABR), which is described in more detail below (section Auditory brainstem response). Subsequently, animals were exposed to a 115-dB SPL narrowband noise (4-kHz center frequency, 10-ms ramps) for 75 minutes, using a free field tweeter (Peerless XT25TG30-04, Quint Audio) placed at 4 cm from the ipsilateral ear. The contralateral ear canal was plugged with a wax mold (Ohropax) and a piece of a foam ear plug on top that filled in the pinna (Herriestoppers). Immediately following noise exposure, ABRs were recorded again to determine noise-induced threshold shifts. Finally, the animal was carefully monitored while it was waking up from anesthesia. Noise exposure and ABR recordings were performed in the same sound-proofed, anechoic chamber as the GPIAS measurements. In 4 out of 19 animals, the noise was not played but all other procedures were carried out. These sham-exposed animals were used as a control group.

### Auditory brainstem response

#### Before and immediately after noise exposure

The auditory brainstem response (ABR) was recorded before and immediately after noise exposure. Platinum Grass needle electrodes (Natus) were placed subdermally on the vertex and caudal-ventrally to the bulla, as reference and active electrodes, respectively. The recorded signal was amplified (10,000 x) and bandpass filtered (300 – 3,000 Hz) by an isolated bio-amplifier (ISO 80, World Precision Instruments [WPI]) and stored on a computer using an external audio card (Fireface UCX, RME Audio; 48 kHz sampling rate).

Acoustic stimuli were presented using the same Fireface audio card (48 kHz sampling rate), pre-amplified using a power amplifier (RMB 1506, Rotel) and presented in free field through a multi-field magnetic speaker (MF1, Tucker-Davis Technologies [TDT]). ABR data collection and acoustic stimulus presentation through the audio card was controlled by custom-written software in MATLAB (Mathworks). ABRs were measured in response to tone bursts of 2, 4, 8, and 16 kHz (10-ms duration, 1-ms on- and offset ramps). Tone bursts were presented at a range of levels that were presented randomly between 5-dB steps. For each frequency, ABRs were collected until a threshold could be visually determined (typically between 200 – 400 repetitions per level). All stimuli were calibrated and equalized using a ½” free-field microphone (type 40AF, GRAS) connected to a pre-amplifier (type 26AK, GRAS) and a power module (type 12AD, GRAS), and custom-written software in MATLAB. ABR traces were revisited offline, and thresholds were visually defined as the lowest sound level at which clear ABR waves were still distinguishable.

#### Preceding single-unit recordings

ABRs were recorded again three days after noise exposure, right before single-unit recordings were collected. The used methods, hardware, and software are similar as described above, with a few exceptions. The reference electrode was placed in the neck muscle, since the necessary surgeries to fix the animal’s skull did not allow for an electrode placement on the vertex. Furthermore, a different audio card was used (Hammerfall DSP Multiface II, RME Audio) and the acoustic stimuli were presented through a headphone driver (HB7, TDT) and a small speaker (IE-800, Sennheiser). The speaker was sealed into an ear bar, which was placed onto the bony ear canal after the pinna was surgically removed (see below at Single-unit auditory nerve fiber recordings – *Surgical preparation*). Stimuli were equalized and calibrated by measuring the sound pressure level near the eardrum with a miniature microphone (ER7-C, Etymotic Research) sealed in the same ear bar, amplified by a microphone amplifier (MA3, TDT). The ABR was measured in response to the same tones as described above with two exceptions. Due to the high-frequency roll-off at 16 kHz for the ER7c microphone, the tone burst at 16 kHz was replaced by a tone burst at 14 kHz. Furthermore, the ABR was also measured during the presentation of chirps (0.3-19 kHz, 4.2-ms duration), to derive a threshold generated by a large portion of the cochlea which could be easily re-evaluated during the course of single-unit recordings. These ABR recordings were carried out in a custom-build sound-attenuating chamber. The overlapping thresholds of the sham-exposed animals between the two setups suggests that, at least in terms of thresholds, these experimental setups are comparable (Fig. 1B).

### Single-unit auditory nerve fiber recordings

#### Surgical preparation

Three days following noise exposure and one day following post-exposure GPIAS evaluation, recordings were made from single AN fibers. Gerbils were anesthetized using the same anesthetic procedures as described for noise exposure (see above at Noise exposure – *Anesthesia*). In this set up, the head of the animal was fixed in a bite bar with a head mount (Kopf Instruments). After removing the pinna, the ear bar with the sound system was placed onto the bony ear canal and sealed with petroleum jelly. The bulla was ventilated through a small, dorsal-lateral hole. Subsequently, the AN was accessed dorsally by making a craniotomy and duratomy over the ipsilateral cerebellum. Cerebellar tissue was partially aspirated, and the AN was exposed by placing a few small saline-drenched parts of latex gutta-percha points (Kent Dental) between the temporal bone and the brainstem.

#### Experimental setup

Activity from single AN fibers was recorded using glass micropipette electrodes (BF120F-10, Science Products) with a typical impedance between 20 and 40 MΩ (pulled with P-2000, Sutter Instruments and filled with 3M KCl solution). Signals from the electrode were amplified (10x, WPI 767), filtered (50/60 Hz; Hum Bug, Quest Scientific), made audible (MS2, TDT), visualized (SDS 1102CNL, SIGLENT Technologies), and digitized (RX6, TDT; 48,828 Hz sampling rate). An Ag/AgCl pellet electrode (Warner Instruments) that was placed subdermally in the neck served as an electrical reference. Acoustic stimuli were presented by a small speaker (IE 800, Sennheiser) that was sealed in the ear bar. Stimuli were calibrated for each animal individually using a miniature microphone (ER7c, Etymotic Research) that was positioned at the exit of the ear bar, close to the tympanic membrane, and custom-written MATLAB code. To isolate single fibers, broadband noise bursts (60 – 75 dB SPL) were presented while slowly advancing the electrode through the auditory nerve using a piezo microdriver and handset (6000 ULN and 6005 ULN handset, Burleigh), until spikes could be detected. Custom-written MATLAB software controlled stimulus presentation and data recording. The single-unit experiments were carried out in a custom-build sound-attenuating chamber.

#### Data recording

After a single unit was isolated, spikes were identified using a simple, adjustable voltage detection threshold. Best frequency (BF) was determined by presenting tone bursts at a range of frequencies (50-ms duration, 5-ms cosine ramps, 5 repetitions, ∼10 dB above the estimated threshold). Next, the fiber’s threshold was determined by presenting tone burst at BF at a range of levels (50-ms duration, 5-ms cosine ramps, 10 repetitions, 3-dB steps). Long recordings in silence, to determine spontaneous rate (SR) and inter-spike interval statistics, were then collected. Each trial was 2.4 s long and a maximum of 75 trials were recorded, resulting in a total of 3 minutes of spontaneous activity. When the fiber was lost during this recording, data was analyzed only for the trials where a good connection to the fiber was ensured. For fibers with a very low SR, the connection with the electrode to the fiber was checked after finishing the recording in silence by presenting tones at BF. When a good connection to the fiber was retained, the following recordings were also obtained. Responses to clicks (two-sample condensation clicks, 97 dB pe [peak equivalent] SPL, 300 repetitions) to determine click latency and responses to pure tones at BF at 30 dB above threshold (50-ms duration, 5-ms cosine ramps, 300 repetitions) to obtain peri-stimulus time histograms.

#### Data analysis

First, threshold for spike detection was revisited offline on a trial-by-trial basis. Next, the frequency-response curve was obtained by plotting the mean response rate as a function of tone frequency. A smoothing spline function was fitted to the curve, and the BF was determined as the peak of this function. When two peaks were observed, which is a known effect of noise exposure on auditory nerve fiber tuning [40, 56], the peak with the highest frequency was considered BF. The rate-level function (RLF) was obtained by plotting mean response rate as a function of tone level. Threshold was determined by the level at which the mean rate was above SR + 1.2*STD of SR and above SR + 15 spikes/s. For this calculation, mean SR and its STD were determined from 10 repetitions of 80-ms silent trials collected at random times during the RLF recording.

Spontaneous rate was determined from the 3-min long recordings in silence. When this recording was not available, the spontaneous rate was estimated from the silent trials of the RLF recording. In addition, the long recordings in silence were used to investigate the statistics of the spontaneous activity. From the inter-spike intervals, which is the invers of the instantaneous spontaneous rate, the coefficient of variation (CV) was calculated. In addition, bursting activity was assessed by applying the robust Gaussian surprise method, according to Ko, Wilson (31), with a minimum number of 3 spikes being considered a burst. Burst rate, number of spikes per burst, and proportion of spikes that were in a burst, were calculated for each recording.

When responses to clicks were available, the click latency was calculated as the first incidence when two consecutive bins (0.05-ms bin size) in the PSTH exceeded the highest bin before the click onset, according to Köppl (57). Furthermore, the mean and variance of the first-spike latency in response to clicks were calculated.

All units included in this study were checked for characteristics typical of auditory nerve fibers. These were: 1) the absence of a pre-potential to the spike waveform, 2) a flat-saturating, sloping-saturating, or straight shape of the rate-level function, 3) a click latency conform with the latency over BF distribution from normal-hearing young-adult gerbils [29], and 4) a primary-like response to tones at 20 – 30 dB above threshold, as described in detail in Heeringa (58).

### Statistical analysis

Following *PPI* calculation, data of the baseline sessions were combined. Data was logarithmically transformed to establish a normal distribution of the data to allow for parametrical testing. *PPI* values were statistically compared between baseline and post-noise exposure sessions for each center frequency of the background noise for each individual animal. Two-sample t-tests, Bonferroni corrected for multiple comparisons, were used to determine significant differences between baseline and post-noise exposure sessions. When *PPI* was significantly increased after noise exposure (worse gap detection) for any for the center frequencies, the animal was labeled with behavioral signs of tinnitus, otherwise the animal was labeled with no behavioral signs of tinnitus. For brevity in the graphs, animals with behavioral signs of tinnitus will be termed ‘TI’, whereas animals with no such behavioral signs in the GPIAS paradigm will be termed ‘NT’.

To determine statistical significance between the experimental groups, one of the following statistical tests was used, wherever appropriate: the Kruskal-Wallis test, followed by post-hoc Wilcoxon Rank sum tests, two-way analysis of variance (ANOVA), followed by post-hoc one-sample or two-sample t-tests, and analysis of covariance (ANCOVA). The Bonferroni-Holm correction was applied whenever a correction for multiple comparisons was appropriate. These statistical analyses were carried out in MATLAB using the Statistics and Machine Learning Toolbox (v23.2). A bootstrap analysis was conducted to examine significance of a change in distribution of spontaneous rate classes between sham- and noise-exposed groups using custom-written MATLAB scripts.

## Acknowledgements

I would like to thank Holger Schulze, Konstantin Tziridis, Richard Gerum, and Rainer Beutelmann for their help in installing and validating the GPIAS experimental setup, and Imme IJsseldijk and Moustafa Almanla for their help in data collection and analysis. Furthermore, I am grateful to Sonja Pyott for her valuable comments on the manuscript.

